# BigMPI4py: Python module for parallelization of Big Data objects

**DOI:** 10.1101/517441

**Authors:** Alex M. Ascension, Marcos J. Araúzo-Bravo

## Abstract

Big Data analysis is a discipline with a growing number of areas where huge amounts of data is extracted and analyzed. Parallelization in Python integrates Message Passing Interface via mpi4py module. Since mpi4py does not support parallelization of objects greater than 2^31^ bytes, we developed BigMPI4py, a Python module that wraps mpi4py, supporting object sizes beyond this boundary. BigMPI4py automatically determines the optimal object distribution strategy, and also uses vectorized methods, achieving higher parallelization efficiency. BigMPI4py facilitates the implementation of Python for Big Data applications in multicore workstations and HPC systems. We validated BigMPI4py on whole genome bisulfite sequencing (WGBS) DNA methylation ENCODE data of 59 samples from 27 human tissues. We categorized them on the three germ layers and developed a parallel implementation of the Kruskall-Wallis test to find CpGs with differential methylation across germ layers. We observed a differentiation of the germ layers, and a set of hypermethylated genes in ectoderm and mesoderm-related tissues, and another set in endoderm-related tissues. The parallel evaluation of the significance of 55 million CpG achieved a 22x speedup with 25 cores. BigMPI4py is available at https://gitlab.com/alexmascension/bigmpi4py and the Jupyter Notebook with WGBS analysis at https://gitlab.com/alexmascension/wgbs-analysis

## 1 Introduction

The term Big Data has been applied since the 1990s [1], and has gained popularity since 2010 due to the exponential amount of data from diverse sources. Until 2007, the volume of all available data was estimated to be slightly less than 300 EB [2]; but in 2018, more than 2.5 EB of data were generated daily [3] due to the vast number of users of social networks, online business, and other platforms. Big Data applications are also used in many branches of science such as physics, astronomy, omics (like genomics or epigenomics [4] [5] [6]). Omics research is currently tightly bound to the rising number of patient records in medical sciences and the application of Next Generation Sequencing (NGS) technologies ([7]). The cumulative amount of genomic datasets in the SRA (Sequence Read Archive) database increased to 9000 TB in 2017 [8] and to over 13000 TB in 2020 [9].

Big Data is defined as a consequence of three shifts in the way information is analyzed [10]: (1) All raw data is analyzed instead of a sample. (2) Data is usually inaccurate or incomplete. (3) General trends and correlations overtake analysis of focused characteristics and minute details.

There are multiple programming languages that can deal with Big Data. Among them, Python is commonly used since it is suited for beginner and experienced programmers; it is dynamically typed and considerable time is saved in code production; and it integrates a vast amount of modules for data analysis, such as pandas, numpy, scipy or scikit-learn, partially implemented in C to optimize computation times.

The requirements of Big Data have prompted the development of a set of tools for data processing and storage such as Hadoop [11], Spark, NoSQL databases, or Hierarchical Data Format (HDF) [12]. Spark is gaining popularity in Python with PySpark module, and there are other libraries like Dask, that implement routines for parallelization using common modules like numpy, pandas or scikit-learn, suiting them to apply Machine Learning techniques to Big Data. Furthermore, there are some recent parallelization libraries, like ray [13], which seem promising for management of large amounts of data in the near future.

One disadvantage of many of the previous libraries is that they do not fully implement all functions from the common Python modules (e.g. Dask does not implement many functions from pandas or numpy), or are limited to adapt to complex algorithms which require extra modules, making them not fully suitable for complex software development. MPI is still a widely used communication protocol for parallelization, more specifically suited for HPC, and Open MPI is one of the most commonly applied implementations of MPI, since it is open source and constantly updated by an active community [14]. MPI4py [15] is the most used module that allows the application of MPI parallelization on Python syntax, allowing users to avoid adapting their pipelines to C++ language, supported by MPI, decreasing code production time.

Nonetheless, there is a limitation in MPI which impedes parallelization of data chunks with more than 2^31^ = 2147483648 elements [16] [17] [18]. The MPI standard uses C int for element count, which has a maximum value of 2^31^ for positive values and, thus, any object with more elements than that cannot be communicated by MPI. This limit is indeed an upper bound since many Python objects, e.g. numpy arrays or pandas dataframes, have a complex structure in which one element of the object corresponds to several C-type elements, decreasing that upper bound to around 10^6^ to 2·10^7^ elements. Communication of bigger objects to the cores throws an OverflowError. The problem also arises with an object small enough for distribution to the cores which, after computation, is transformed into an object too large to be recovered to the original core, throwing an OverflowError. Thus, the size limitation problem negatively affects algorithms whose final object has a undetermined object size, hampering to know in advance whether the size limitation will be fulfilled.

Several solutions to the size limit problem were proposed. Hammond et al. [18] [19] developed BigMPI, that circumvents the C int problem using derived C types allowing up to ~ 9 · 10^9^ GB of data for parallelization. However, BigMPI developers acknowledge the limitation of BigMPI of supporting more complex datatypes, which must be derived to built-in types [19], posing a problem for users whose pipelines include Python-like objects, which would need to be transformed into C-type objects for parallelization. Another solution is to divide the parallelizable object into chunks, and parallelize each chunk independently. The main drawback of this method, implementable in Python, is that it has to be tailored to suit the object type, i.e., a pandas dataframe and a numpy array have to be divided using different syntax. Moreover, the number of chunks that have to be produced to parallelize without error is not straightforward to calculate.

In order to overcome this last problem we developed BigMPI4py. BigMPI4py is a Python implementation which uses MPI4py as the base parallelization module, and applies an optimal object splitting strategy to automatically distribute large objects across processors. BigMPI4py distributes the most common Python data types (pandas dataframes, numpy arrays, simple lists, nested lists, or lists composed of previous elements), regardless of the size, and keeping an easy syntax. BigMPI4py currently allows collective communication methods bcast(), scatter(), gather(), or allgather(), as well as the point-to-point communication method sendrecv(), adapted from the homonym functions on MPI4py. BigMPI4py also communicates numpy arrays with vectorized Scatterv() and Gatherv() methods, providing a substantial time optimization.

The analysis of whole genome scale DNA methylation data is a computational demanding task in omics data analysis since biologically relevant DNA methylation information can be influenced by a single CpG locus and have important regulatory roles in all steps of the life from developmental biology [20], [21], to aging [22] and multiple diseases [23], [24].

In order to identify statistically significant differentially methylated CpG loci across tissues from the three germ layers we have implemented a parallelized version of the Kruskall Wallis (KW) test using BigMPI4py. We have applied this implementation on whole genome bisulfite sequencing (WGBS) 59 samples from 27 different tissues from the ENCODE project [25] to obtain relevant genes with differential methylation across germ layers.

## 2 System overview

BigMPI4py has 5 functions which mimic the actions of MPI and MPI4py: bcast(), scatter(), gather() and allgather() belong to the collective communication category, and sendrecv() belongs to the point-to-point communication category.

- bcast() communicates replicates of an object from the source (root) core to the remaining cores.
- scatter() splits the object from the source core into *n* chunks (*n* = number of cores) and distributes them to the remaining cores.
- gather() combines the objects from all the cores into a unified object and sends it to a destination core.
- allgather() combines the objects from all the cores and distributes copies of the combined object to all the cores.
- sendrecv() communicates an object to a specific destination core.

Due to similarities in the algorithmic structure of these 5 functions, they are combined into 2 unified functions: _general_scatter() and _general_gather(). _general_scatter() is called by bcast(), scatter() and sendrecv(), whereas _general_gather() is called by gather() and allgather(). _general_scatter() has 3 main arguments:

- object type: pandas dataframe, series, numpy arrays or lists. Lists are divided into “simple” lists when they contain integers, floats or strings; and “complex” lists when they contain dataframe, series, arrays or other lists. “Mixed” lists with “complex” and “simple” types of elements simultaneously are not currently supported.
- optimize: if True and when the object is a numeric numpy array it can be scattered using the comm.Scatterv() function from MPI4py. This function uses a vectorized implementation of the parallelization, drastically improving parallelization efficiency.
- by: columns referring to one or several categorical variables. E.g., if a table contains one column with months and another one with week days, choosing by with these two columns selects all combinations of months and week days and distributes rows of the table so that all rows with the same week day and month are distributed to the same core.

_general_gather() arguments are object type and optimize.

Both functions follow these steps:

1. Determine the object type. Objects of different types require different partitioning strategies.
2. Determine the positions to divide the object into *n* parts, or chunks, *n* being the number of processors.
3. Determine, for each chunk of the object, secondary parameters to further divide this object in case of memory limitations.
4. Perform the division of the object.
5. Merge all the communicated objects into a final object.

During the step of determining the positions, a list of indexes is created whereby the object *A* is split into *n* chunks. *A* will be divided into equally-sized chunks unless by argument is set; in that case the number of combinations will be equally distributed.

If *A* has length |*A*| (number of rows in arrays, dataframes or series, and number of elements in lists), then the index positions of the division are

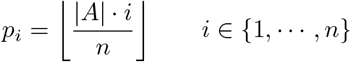

*A_i_* is defined as the *i*-th chunk of *A* and, the *i*-th chunk would start at position *p*_*i*–1_ and end at position *p_i_*. If by argument is set to True, then *p*_*i*–1_ and *p_i_* would be the indexes that separate the two categories.

It is possible that some *A_i_* chunk exceeds the maximum size, and the object cannot be parallelized. Thus, during the step of determining the positions, instead of dividing the object into *n* chunks, each of the chunks *A_i_* is further split into k subchunks, so that the object size restriction is overcome during the communication of *A* to the processors.

To calculate k, two strategies are developed, hereby termed Strategy I and Strategy II. Those strategies are slightly different depending on whether an scattering or a gathering is being performed, since the final object will be distributed to all the cores, or will be communicated to a specific one.

### 2.1 Strategies to calculate the number of subchunks

#### 2.1.1 Strategy I

Strategy I deals with “simple” and “complex” lists, arrays, dataframes and series. The size (memory allocation) of *A_i_* is designated as 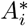. After *A* has been divided into *A*_1_, *A*_2_, …, *A_n_*, if 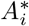 is greater than *L_n_* = *L/n*, *L* being the memory limit (which can be assigned by the user), then *k_I_* is defined as

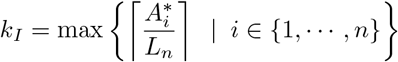

BigMPIpy defines *A_i,k_i__* as the *k_i_*-th subchunk of the *i*-th chunk. By choosing this *k_I_* value, it is assured that any combination (*i*, *k_i_*) will result in an *A_i,k_i__* smaller than *L_n_*. Therefore, *A* could be expressed as the following matrix of subchunks:

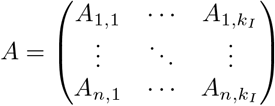

The positions by which the objects are divided are:

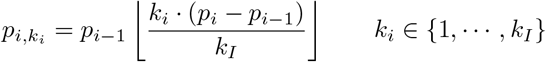

For each value *k_i_*, all *A_i,k_i__* are combined into a list, and communicated by MPI4py functions. The model of organization varies depending on whether the object is scattered or gathered, although the main strategy remains the same. The description of the scattering and gathering algorithms are shown in Algorithms S1 and S2 respectively.

#### 2.1.2 Example of array scattering with Strategy I

In this case we will distribute an object *A* into *n* = 5 cores as illustrated in Fig. 1 where the object *A* is represented with a pentagon. Since *n* = 5 *A* is divided into 5 chunks: *A*_1_, *A*_2_, *A*_3_, *A*_4_, and *A*_5_, represented with triangles. If *L* = 3, then *L_n_* = 3/5 = 0.6. if the weights of the chunks are 1, 1.5, 1.2, 1.1 and 1 respectively, *k_I_* = ⌈1.5/0.6⌉ = 3.

**Fig. 1.**
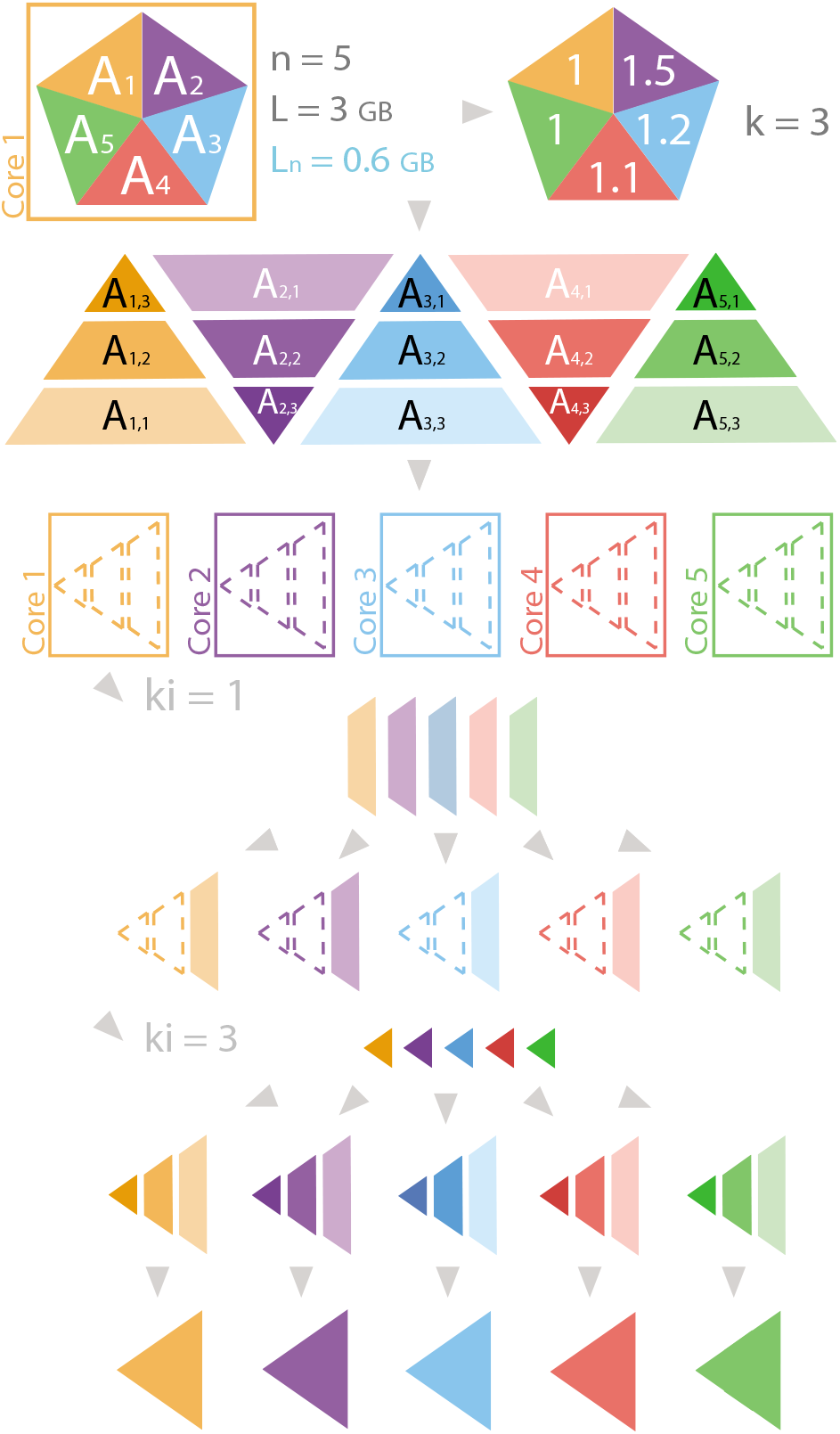
Example of array scattering with Strategy I. *n* is the number of cores, *L* is the memory limit (in GB) and *L_n_* = *L/n*. For each piece, the hue represents the core that will process the chunk or subchunk, and the luminosity represents the *k_i_* loop in which the subchunk will be processed. Joined pieces belong to the same object, whereas separated pieces represent a list with subchunks. Dashed lines represent Nonetype objects within a list.

Then, the distribution of subchunks is *A*_1,1_, *A*_1,2_, *A*_1,3_, *A*_2,1_, ⋯ …, *A*_5,2_, *A*_5,3_, which are combined in the list scatter_A. During the first iteration on *k_I_*, all *A*_*i*,1_ are combined, distributed and attached to scatter_object, i.e., the third core receives *A*_1,3_. After all iterations, each core has the list scatter_object = [*A*_*i*,1_, *A*_*i*,2_, *A*_*i*,3_], which, after merging, will be transformed to *A_i_*.

#### 2.1.3 Example of array gathering with Strategy I

Let’s suppose that *n* = 3 and there are three arrays to be gathered: *A*_1_, *A*_2_, *A*_3_; as illustrated in Fig. 2, where each array is represented with a rhomboid and the final gathered array with a triangle. If *L_n_* = 2 and 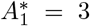, 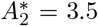, 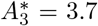, then *k* = ⌈max{3/2, 3.5/2, 4/2}⌉ = 2. Therefore, *A_i_* is split into *A*_*i*,1_ and *A*_*i*,2_. In the receiver core gather_object is set to [None, None, None, None, None, None]; and after the first iteration on *k_I_*, the receiver core receives gather_ki= [*A*_1,1_, *A*_2,1_, *A*_3,1_], and gather_object is [*A*_1,1_, None, *A*_2,1_, None, *A*_3,1_, None]. After the second iteration, gather_object is [*A*_1,1_, *A*_1,2_, *A*_2,1_, *A*_2,2_, *A*_3,1_, *A*_3,2_], which is merged onto the object *A* represented with a triangle in Fig. 2. Gathering a list with Strategy I involves similar steps as gathering an array.

An example of a list scattering with Strategy I can be found at Supporting Information.

**Fig. 2.**
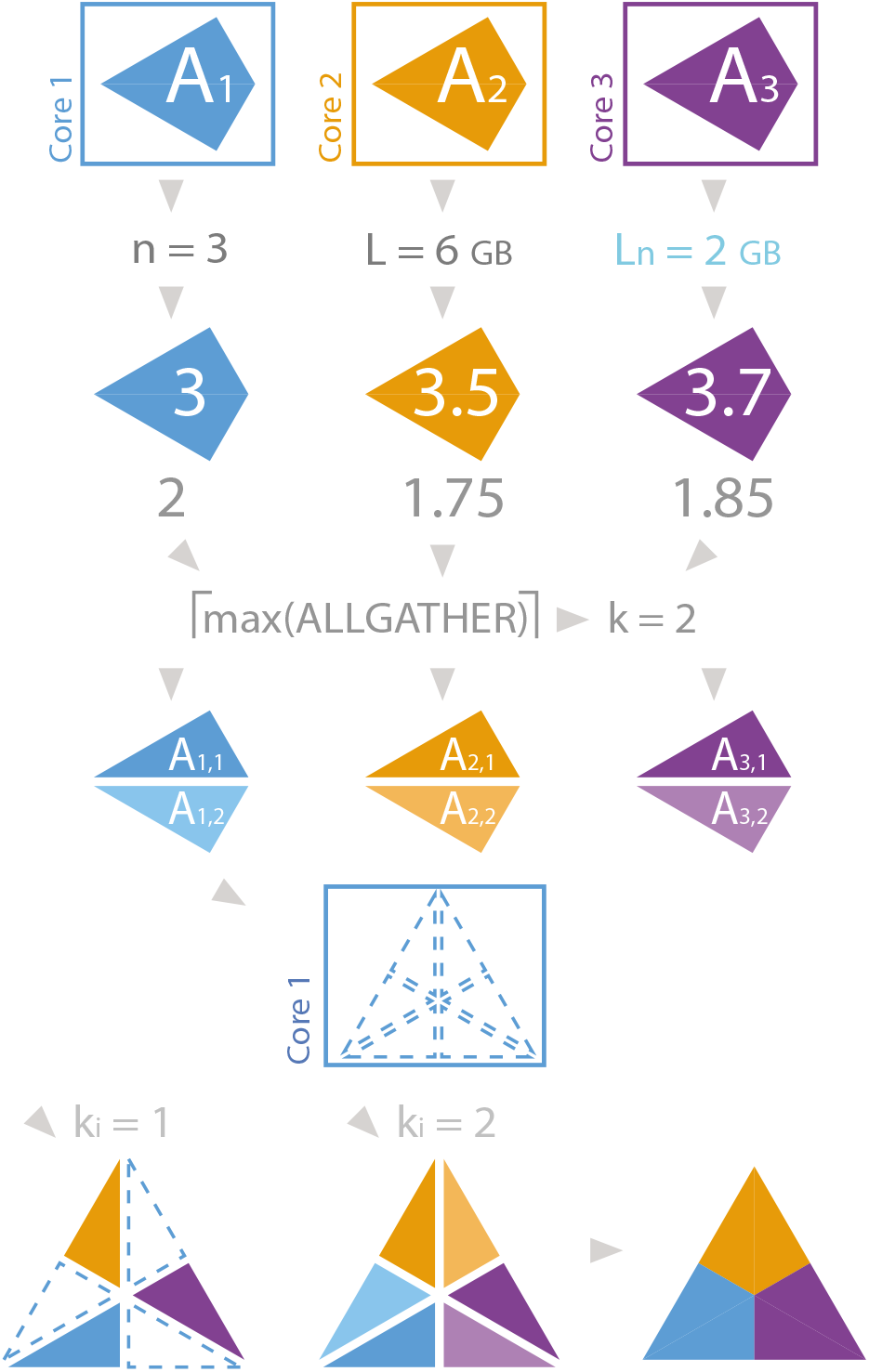
Example of array gathering with strategy I. *n* is the number of cores, *L* is the memory limit (in GB) and *L_n_* = *L/n*. For each piece, hue represents the core that will process the chunk or subchunk, and luminosity represents the *k_i_* loop in which the subchunk will be processed. Joined pieces belong to the same object, whereas separated pieces represent a list with subchunks. Dashed lines represent Nonetype objects within a list.

#### 2.1.4 Strategy II

Strategy II deals with “complex” lists *A* in which their elements *a_i_* can be partitioned, and one or more of them individually exceeds the memory limit.

In this case, the *k_I_* value from strategy I is not suitable since the best partition of elements would be the one that makes each element of the list to be scattered or gathered individually. E.g., for *A* = [*a*_1_, *a*_2_, ⋯, *a*_4_, *a*_5_], with the memory allocation required for 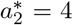 units of memory and *L_n_* = 2, if *n* = 2, then the *k* value for this case would be *k_I_* = 3, making each element to be scattered individually. However, since 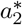 alone is greater than *L_n_*, the scattering would be impossible.

Given a list *A* with *m* elements *A* = [*a*_1_, ⋯, *a_m_*], the main objective of Strategy II is to split each element of *A* into *k_II_* sub-elements *a_i,k_i__*, so that the sum of the lengths of all *a_i,k_i__* elements, for a fixed *k_i_*, is smaller than *L*. Those elements will then be communicated and gathered into respective lists for each core. Thus, *k_II_* is defined as

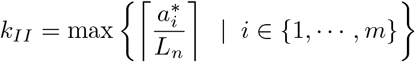

where *n* is the number of cores and *m* is the number of elements of the list.

The description of the scattering and gathering algorithms are shown in Algorithms S3 and S4 respectively.

#### 2.1.5 Example of scattering with Strategy II

Assume that we want to distribute an object *A* = [*a*_1_, *a*_2_, *a*_3_, *a*_4_, *a*_5_, *a*_6_] into *n* = 2 cores, and *k_II_* = 3, as represented in Fig. 3. The distribution list is [[*a*_1_, *a*_2_, *a*_3_], [*a*_4_, *a*_5_, *a*_6_]]. Then, each element *a_i_* is split into [*a*_*i*,1_, *a*_*i*,2_, *a*_*i*,3_], and the previous list becomes scatter_list = [[[*a*_1,1_, *a*_1,2_, *a*_1,3_], [*a*_2,1_, *a*_2,2_, *a*_2,3_], [*a*_3,1_, *a*_3,2_, *a*_3,3_]], [[*a*_4,1_, *a*_4,2_, *a*_4,3_], [*a*_5,1_, *a*_5,2_, *a*_5,3_], [*a*_6,1_, *a*_6,2_, *a*_6,3_]]]. For each *k_i_* (in this example, *k_i_* = 1) the list merge_ki is created: [[*a*_1,1_, *a*_2,1_, *a*_3,1_], [*a*_4,1_,*a*_5,1_,*a*_6,1_]]. After the scattering, the first core receives [*a*_1,1_, *a*_2,1_, *a*_3,1_] while the second core receives [*a*_4,1_, *a*_5,1_, *a*_6,1_]. This list is appended to the list merge. After all *k_i_* values, merge is [[*a*_1,1_, *a*_2,1_, *a*_3,1_], ⋯, [*a*_1,3_, *a*_2,3_, *a*_3,3_]] for core 1 and [[*a*_4,1_, *a*_5,1_, *a*_6,1_], ⋯, [*a*_4,3_, *a*_5,3_, *a*_6,3_]] for core 2. Then, [*a*_*i*,1_, *a*_*i*,2_, *a*_*i*,3_] is merged to *a_i_* for *i* = 1, ⋯, 6, and return_list is [*a*_1_, *a*_2_, *a*_3_] for core 1 and [*a*_4_, *a*_5_, *a*_6_] for core 2.

**Fig. 3.**
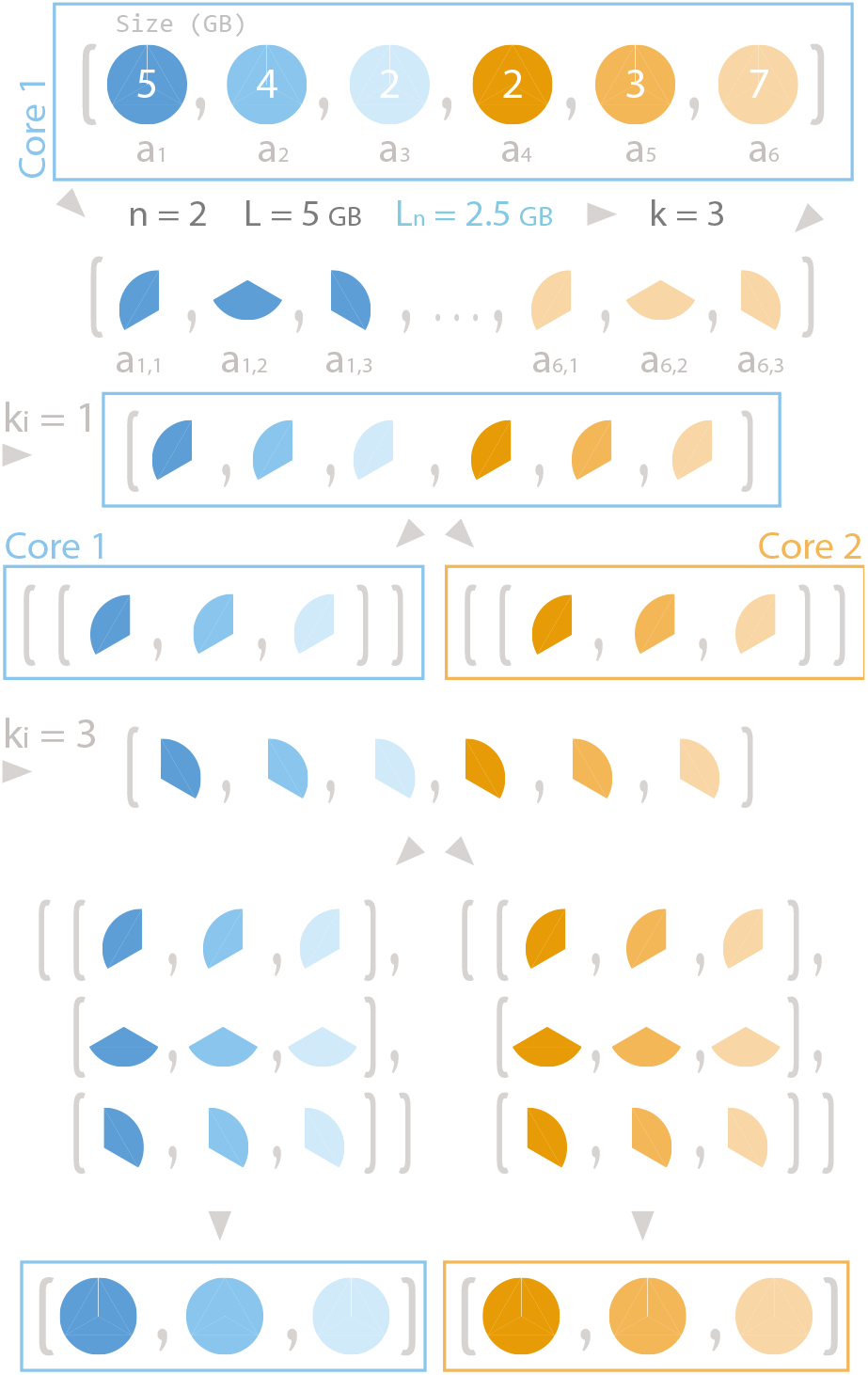
Example of array scattering with strategy II. *n* is the number of cores, *L* is the memory limit (in GB) and *L_n_* = *L/n*. For each piece, the hue represents the core that will process the chunk or subchunk, and the luminosity represents the *k_i_* loop in which the subchunk will be processed.

An example of a gathering with Strategy II can be found at Supporting Information.

### 2.2 sendrecv(), allgather() and bcast() functions

sendrecv() function was implemented for point-to-point communication, whereas allgather() and bcast() were implemented for collective communication of whole objects across cores. sendrecv() and scatter() share Strategy I and Strategy II, and were together in _general_scatter(). The main differences in the use of scatter() and sendrecv() are:

1. sendrecv() sends the object to a single core, thus *n* = 1 is set.
2. scatter() calls comm.scatter() for scattering, whereas sendrecv() calls comm.send(object, dest) (where dest is the destination core) and comm.recv(root) for communication of the object.
3. Merging of objects with Strategy II in sendrecv() is simplified since *n* = 1.

allgather() was integrated together with gather() in _general_gather(), since Strategy I and Strategy II are also shared. The main differences in the use of allgather() and gather() are:

1. Instead of calling comm.gather(), comm.allgather() is called.
2. Combination of subchunks is performed for all cores instead of the destination core only.

bcast() function was implemented in a separate function, with the strategy developed in Algorithm S5. If the available memory of the computer is smaller than *n*·*A*^*^, a MemoryError is thrown before *A* is broadcasted.

### 2.3 Vectorized implementation of gather() and scatter()

MPI4py includes vectorized functions Scatterv() and Gatherv() for numeric numpy arrays, and improves communication by transferring the buffer of the chunks, instead of the values of the chunks. BigMPI4py automatically checks whether the object is a numeric array and scatters or gathers the object using the corresponding function. During the vectorized implementation the array splitting positions *p*, the number of rows, and the number of cells in each chunk are calculated, and these values are used with Scatterv() and Gatherv() methods from MPI4py. BigMPI4py adapts the vectorization also for large arrays, supporting values of *k_I_* greater than 1. For this case, when using scatterv(), the list of scattered objects for each *k_i_* is merged into an array, and the counts for each array are considered. Although this step might be computationally expensive for big arrays, time performance is still improved, as shown in Results section.

## 3 Results

### 3.1 Computation time analysis on random data

Scattering and gathering performance was compared using random datasets to avoid apparent faster times due to use of cache memory. The results of the times required for scattering and gathering are shown in Figs. 4 and S1, respectively.

**Fig. 4.**
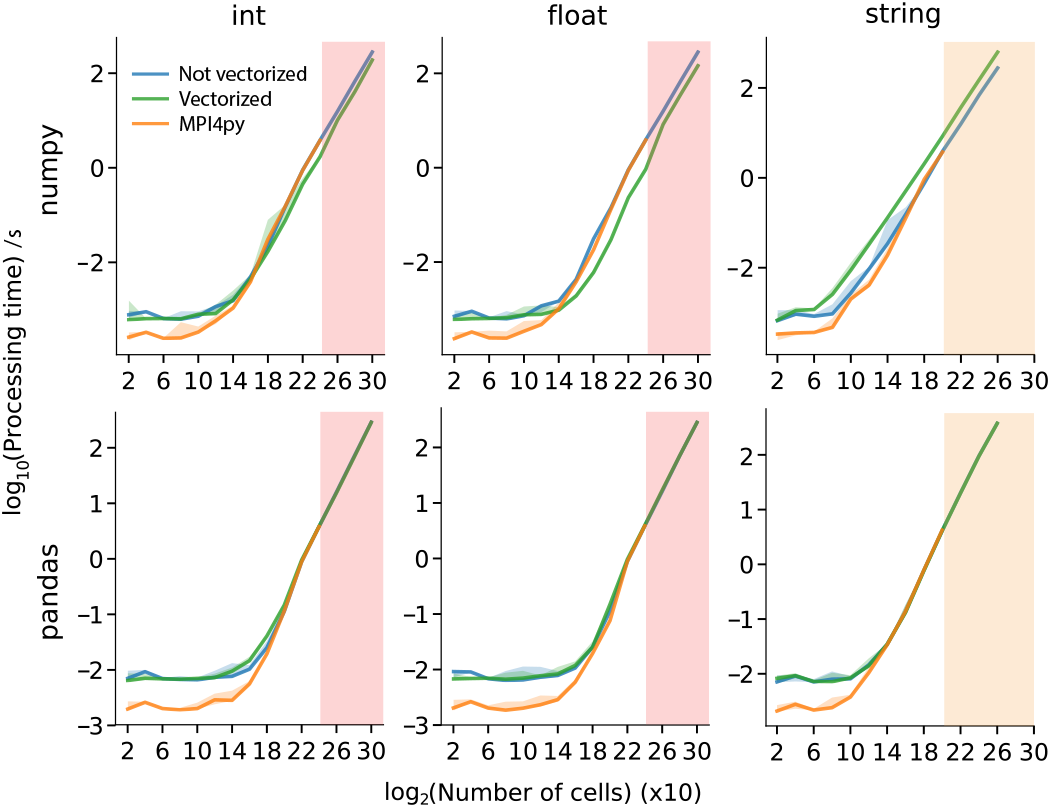
Scattering times for numpy arrays and pandas dataframes for three data types. The number of cells that MPI4py is not able to process due to OverflowError is shadowed in pink for int and float data types, and in orange for string data type. Color filling between lines represents the 10^th^ and 90^th^ percentiles of the data.

Both vectorized and non-vectorized approaches process larger tables than the MPI4py approach. When scattering, int and float tables MPI4py can only process up to nearly 2^24^·10 ≈ 167 million cells, and string tables up to 2^20^· 10 ≈10 million cells; whereas BigMPI4py reaches table sizes up to the maximum available memory. The objects that MPI4py cannot process due to its size limitation have sizes that are common for objects required to be processed in multiple Big Data projects.

Scattering and gathering processing times show a common trend across all tables and types: there is a quasilinear behavior until 2^13^ · 10 ≈ 80000 to 2^15^ · 10 ≈ 300000 cells, in which the processing times are around 10^−3^s for numpy arrays, and 10^−2^s for pandas dataframes. For larger numbers of cells, however, the times increase exponentially (linearly after log-log transformation), following a trend with almost no deviation from the expected regression. For short processing times, MPI4py shows faster computation performance, up to an order of magnitude. This difference is due to the preprocessing done by BigMPI4py, where the main delays are the communication of intermediate objects with MPI4py, like *k_I_* values calculation, consuming up to 60% of the total time. It should be noted that the difference of times in this range is of less than a tenth of a second, and are not relevant for most of the parallelization cases. For longer processing times, most of the time (> 95%) is dedicated to the communication of the main object across cores, or the merging of a list of tables into a single table. For vectorized processing merging can take up to 70% of the processing time, whereas with non-vectorized processing merging takes up to 35% of the processing time. Interestingly, vectorized parallelization is still nearly 10 times faster, even considering that long time dedicated to merging. For string arrays the processing times are longer for the vectorized processing since a conversion between string and float must be performed for vectorized parallelization using Scatterv(), which consumes up to 80% of the total processing time. By default, string arrays are not processed by vectorized parallelization, which should be used for numeric arrays.

Vectorized parallelization is only available for arrays, and not for pandas dataframes, due to the large amount of time that requires to transform a numpy array to a pandas dataframe in case of big object sizes. Thus, processing times for vectorized and non-vectorized parallelization in pandas arrays are technically the same (Figs. S2 and S3).

Numpy float arrays show a great parallelization advantage (up to five times for 2^18^ rows) by vectorization both in scattering and gathering. For numeric types, vectorized scattering shows at least a two-fold time decrease even for number of rows that are not supported by MPI4py due to size limitations. Regarding vectorized gathering, numpy float arrays show improved computation times across the entire size spectrum, although for int arrays the processing time increases in relation to the non-vectorized processing. Profiling shows that >95% of time is associated to barrier synchronization, which implies a mismatch in processing rhythms across cores. This time profiling is highly dependent on the number of cores, obtaining better times with fewer processors, and cannot be always reproduced.

### 3.2 BigMPI4py evaluates the CpG loci methylation statistical significance of WGBS data within optimal parallelization times

In order to test BigMPI4py on the processing of whole genome scale DNA methylation data, we have used the WGBS data of 59 samples of 27 human tissues from the ENCODE project.

As a first task we have classified samples on the three different germ layers (ectoderm, endoderm and mesoderm) based on their CpG methylation status and developed a parallel implementation of the KW test to find CpG loci with statistical significant differential DNA methylation levels across germ layers. A total of 55 million positions were evaluated, from where 27778 CpG loci pass the KW test with a degree of significance of 2 · 10^−6^.

We observe in Fig. 5 that computation times of our parallel implementation of the KW test follow the expected behaviour, that is, doubling the number of CPUs should decrease the processing time in half. The *R*^2^ for the linear fit of the data is 0.9994, which shows that BigMPI4py is able to parallelize the KW test of DNA methylation matrices with times as expected with parallel computing. The parallel implementation achieved a 22x speedup on computation time with 25 cores, allowing the analysis of nearly 55 million CpG methylation loci in less than 20 minutes.

**Fig. 5.**
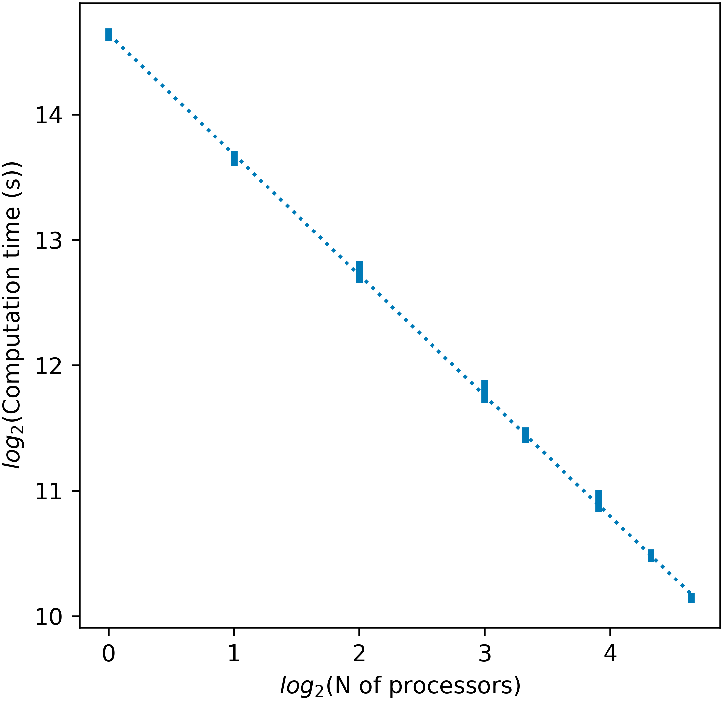
Scattering times for the KW test. Computation time is defined as the difference between the total time, i.e. the time used to run all lines of code, and the baseline time, i.e. the time dedicated to non parallelized actions like writing and reading of data. Baseline computation times are shown in Fig. S4. For each number of processors, vertical bars represent the minimum and maximum computation times, out of 5 trials. Data were fitted to a linear regression, shown in a dotted line.

### 3.3 BigMPI4py allows the detection of genes associated to differentially methylated regions across germ layers

The 27778 CpG loci with a KW test *p*-value smaller than 2 · 10^−6^ show 3 clusters after being clustered by tissue, and 3 main clusters after being clustered by DNA methylation pattern (heatmap in Fig. 6). Tissue classification is almost perfect, with only one replicate of adipose tissue being misclassified as belonging to endoderm.

**Fig. 6.**
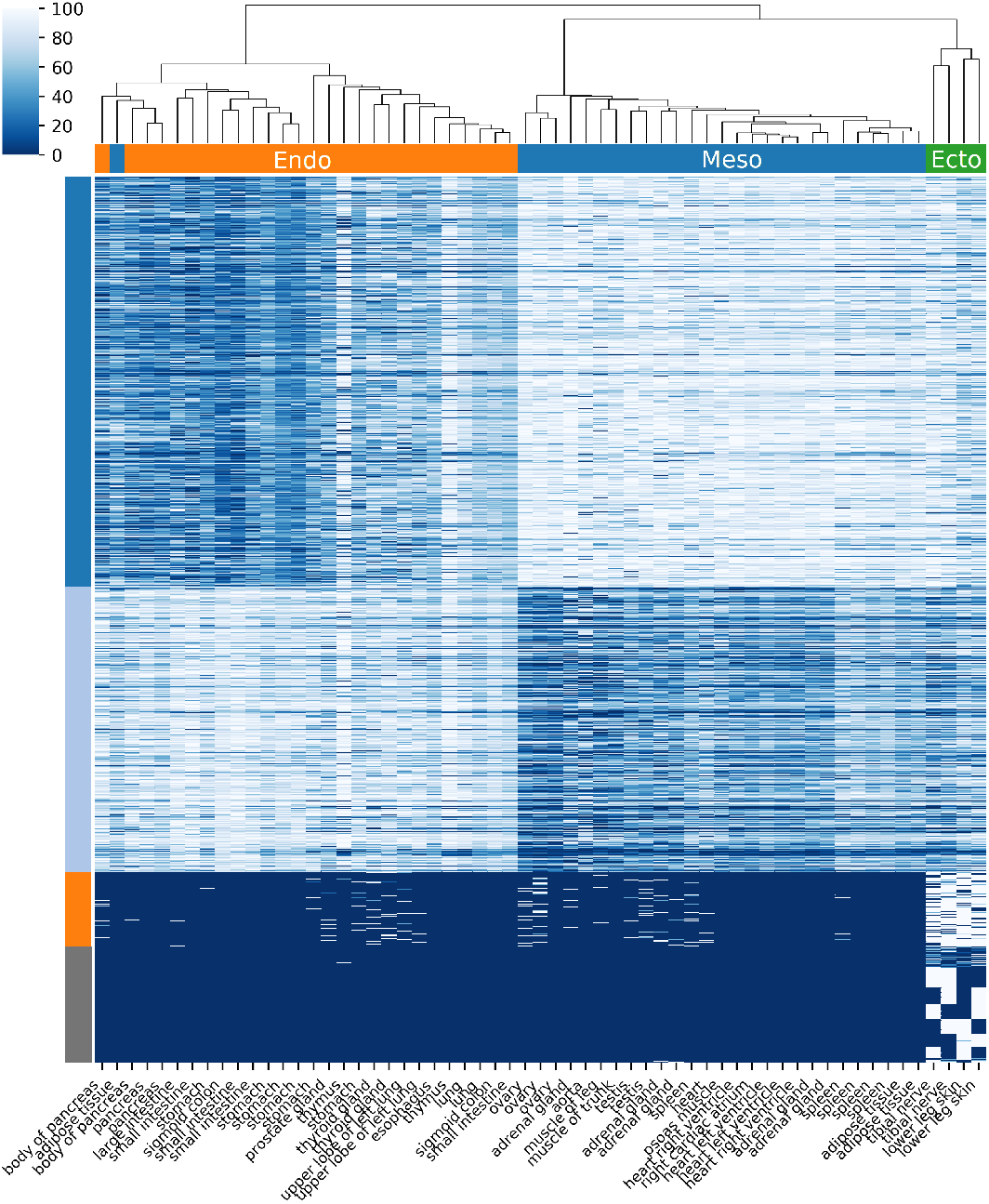
Clustermap of statistical significant differentially DNA methylated CpG loci. Heatmap columns are different manually-labelled tissue types (orange: endoderm, blue: mesoderm, green: ectoderm), and rows are the 27778 CpG loci that passes our KW with a degree of significance 2 · 10^−6^. Colored labels in rows represent the identities obtained applying hierarchical clustering by rows. The color bar in the top left codifies the CpG methylation in percentage. The higher is the methylation the whiter is the color.

Clustering by rows shows 15 clusters, that can be subclassified in 3 different categories: CpG loci with high methylation in ectoderm and mesoderm (topmost cluster in heatmap, dark blue), CpG loci with high methylation in endoderm (second cluster, light blue), and CpG loci with high methylation in ectoderm (third cluster, orange).

After manual inspection of methylation patterns near the genes associated to the CpG loci of the last set of clusters, joined and depictyed in gray, reveals that methylation signal of those statistical significant different methylates CpG loci have not a clear pattern, and can be attributed to random sampling of ectoderm samples, due to low sample numbers in that germ line. This phenomenon is shown for genes NPIPA2 and GATDB3 in Fig. S5. In other words, since the number of tissues associated to ectoderm is much smaller than the number of tissues from the rest of germ layers, finding statistically significant differentially methylated CpG loci with different methylation in one or more of ectoderm samples is much easier than in the rest of germ layers, and the different methylation of those CpG loci can be considered as false positives.

Clustering by rows and columns is robust to *p*-value selection. Higher *p*-values (around 10^−4^) and lower *p*-values (up to 10^−10^, the expected value applying Bonferroni correction) show similar types of clusters, both row- and column-wise. Generally, most of the genes that appear near statistically significant different methylated CpG loci with less stringent *p*-values appear also with more stringent *p*-values.

CpG loci with high methylation in ectoderm and mesoderm (i.e. low methylation in endoderm) can be divided in two subclusters: one associated to gastrointestinal tissues (pancreas, body of pancreas, large intestine, small intestine, colon, and stomach) and the other one associated to the rest of endodermal tissues (thymus, thyroid gland, upper lobe of left lung, and lung), with some gastrointestinal samples from the first subcluster.

The list of selected genes is shown in the first column of Table S1. We observe that most of the associated genes have lower DNA methylation levels on the gastrointestinal tissues (stomach, and some intestine samples), such as BCL2, BHLHE41, DNMBP, ELF3, KLF5, LPCAT3, MIR200B, MIR200CHG, PRR15L, TMEM51, or CDH1, shown in Fig. 7, up. There are no clear associations to gastrointestinal activity in any of those genes, although some of them have been associated to different cancers: CDH1 is associated to gastric, breast, colorrectal, and thyroid cancer [26], [27], [28], [29], KLF5 is associated to colorrectal cancer [30], MIR200B has been shown to promote mesenchymal-to-epithelial transition (MET) in last steps of metastasis [31], MIR200CHG is associated to bladder urothelial carcinoma [32], and PRR15L is associated to sigmoid colon cancer [33].

**Fig. 7.**
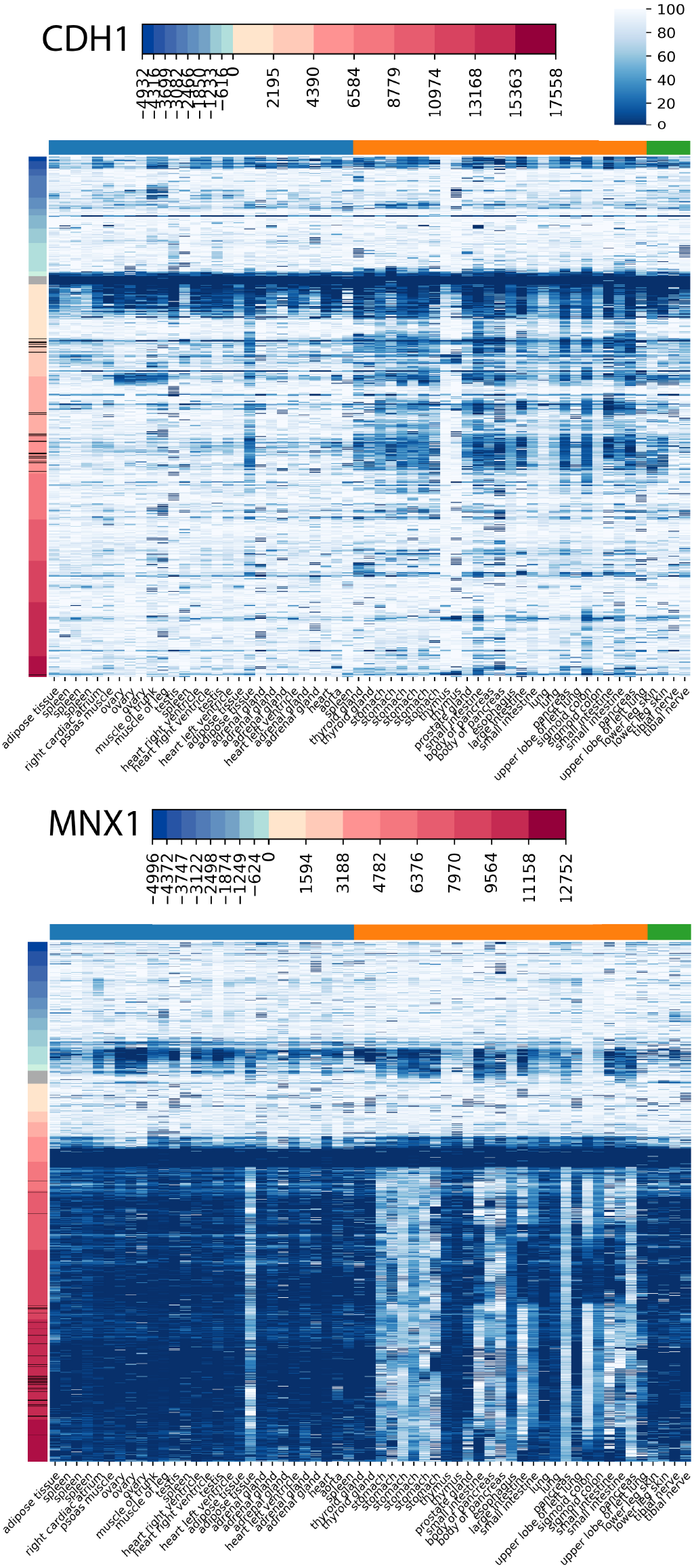
Heatmap of methylation near CDH1 and MNX1. Heatmap columns are different manually-labelled tissue types (orange: endoderm, blue: mesoderm, green: ectoderm), and rows are genomic positions of methylation. Coloured labels in rows represent the distance from the gene, in bases. Gray label represents methylation positions within the gene, blue labels represent positions upstream the gene, and red labels represent positions downstream the gene. Black marks in row labels represent positions with *p*-value < 2 · 10^−6^. The color bar in the top right codifies the CpG methylation in percentage. Higher methylation corresponds to whiter color.

Interestingly, we found in some genes that CpG loci at the same distance to their transcription start sites (TSSs) have uniform methylation levels across most of the tissues but strong reverse signal in others. E.g., there is a region within MIR200CHG gene (Fig. S5) with high DNA methylation in thymus, and slight methylation level in spleen; whereas an upstream region shows a strong demethylation of both tissues, possibly indicating a regulatory activity of this gene in those two immune tissues.

CpG loci with low methylation in ectoderm and mesoderm do not show a distinct pattern of division into subclusters. The list of selected genes for that case is shown in the second column of Table S1. Similar to the former case, selected genes do not always show a clear biological function associated to a germ layer, although some of them have been associated to cancer: FAM198B is associated with inhibition of lung adenocarcinoma [34], IFFO1 is associated to non-small-cell lung cancer [35], LINC00261 is associated to non-small-cell lung cancer and colon cancer suppression [36], [37], MNX1 (Fig. 7, down) is associated to cell proliferation in colorrectal cancer [38], and ST3GAL4 is associated to cervical cancer ‘ [39].

For C1S, CDH1, DDR2, LINC00261, MNX1, PRRT1B, or IFFO1 genes; spleen and thymus have the same level of methylation, which might imply that, despite germ layer being a feature that is distinct for some tissues, there are tissues with a shared function across layers that show that biological function can be the most relevant feature to cluster tissues together.

## 4 Examples of use of BigMPI4py

Besides overcoming the object size limitation of MPI4py, BigMPI4py is designed to follow the philosophy of a simple syntax. Required attributes for BigMPI4py are scatter_object (object to be scattered) and comm (MPI4py.MPI.COMM_WORLD).

To use BigMPI4py the header at Algorithm 1 is required.

### Algorithm 1 Header of a Python file

1 from mpi4py import MPI

2 import BigMPI4py as BM

3 comm = MPI.COMM_WORLD

4 size, rank = comm.Get_size(), comm.Get_rank()

Algorithm 2 shows array, dataframe and complex list scattering and broadcasting.

All variables must be declared as None to do the communication (line 1). Then, for the root core, the object is created (lines 2-5). Finally, scatter() function is called (lines 6-8). The same syntax is applied for object broadcasting (line 9). The code is simple, and with a working knowledge MPI4py, scattering is implemented in 4 lines of code.

If an object is distributed according to some categorical columns, by argument must be used. Algorithm 3 shows an example of this. In this example BigMPI4py extracts all pairs of values from the categorical columns ([‘A’, ‘Red’], [‘B’, ‘Red’], [‘B’, ‘Blue’], [‘A’, ‘Blue’], [‘C’, ‘Red’] and [‘C’, ‘Blue’]), and scatters the dataframe so that no dataframe with each pair of values is distributed across cores.

### Algorithm 2 Object scattering and broadcasting

~~~
1 arr, df, lst = None, None, None
2 if rank == 0:
3    arr = np.random.rand(2 ** 26, 10)
4    df = pd.DataFrame(arr)
5    lst = [arr, df, arr, df, arr, df]

6 scatter_arr = BM.scatter(arr, comm)
7 scatter_df = BM.scatter(df, comm)
8 scatter_lst = BM.scatter(lst, comm)

9 bcast_lst = BM.bcast(lst, comm)
~~~

### Algorithm 3 Object scattering with categorical variables

1 arr = None

2 if rank == 0:

3 arr = pd.DataFrame(

4 [[‘A’, ‘Red’, 0],

5 [‘A’, ‘Red’, 1],

The procedure to code object gathering and allgathering is similar to the procedure of scattering. Algorithm S6 shows the communication of an array for gather and allgather. The rest of object types follow the same syntax. Gathering and allgathering of an object requires only two lines of code, making it even easier to program. An example of array point-to-point communication is shown in Algorithm S7. Any code that uses MPI4py and BigMPI4py must be run externally using an MPI implementation, since it depends on MPI. Algorithm S8 shows an example using mpirun program from OpenMPI implementation.

## 5 Conclusions

BigMPI4py brings the possibility to take advantage of the parallelization implementing MPI in Python without any theoretical object size limitation, using all the computational power (number of cores and memory) of hardware (multicore PC, workstation or HPC) in Big Data projects with the only limitation being the resources in the system. The use of BigMPI4py increases the “robustness” of MPI4py, since it allows to overcome the OverflowError arising in MPI4py when the size of the output object exceeds the limit of MPI4py even when the size of the input object does not exceed such limit. Additionally, BigMPI4py simplifies the use of MPI4py by automatically splitting the object to be communicated across the cores and by automatically deciding whether to optimize the parallelization by performing vectorization of objects.

We have developed a parallelization of the KW test that allow us to find statistically significant different methylated CpG loci across all the WGBS human samples of the ENCODE project.

We found that while endoderm and ectoderm samples have specific high methylated CpG loci, there are no mesoderm-specific high methylated CpG loci, i.e. the mesoderm samples share their high methylated CpG loci with the either ectoderm or endoderm samples. The same occurs with demethylated loci, which are shared between either mesoderm and endoderm, or mesoderm and ectoderm.

Although most of the genes with low methylation levels on the gastrointestinal tissues have no clear associations to gastrointestinal activity, some of them have been associated to different cancers. However, these observations should be taken with a grain of salt considering the bias towards characterization of biological functions as functions observed in cancer.

There are tissues with a shared function of genes across layers showing that biological function could be the most relevant feature to cluster tissues at DNA methylation level together that the germ layer of origin. We found sets CpG loci whose methylation status is very specific for some tissues, flipping strongly for the rest. These loci are candidates to be specific epigenetic regulators of tissue identity.

## 6 Methods

### 6.1 Computation time analysis of random data

To test the time performance of scatter() and gather() functions, we performed several simulations on randomized datasets. For each function, random numpy arrays or pandas dataframes with 10 columns and row numbers ranging from 2^2^ to 2^26^ (gather()) or 2^30^ (scatter()) at step 2 were generated. Different data types (int, float and string) were used during random array generation.

For each case, three functions are compared: the one from MPI4py, and the equivalent ones, vectorized and non-vectorized, from BigMPI4py. All measurements were repeated 5 times.

cProfile and line_profiler Python modules were used for monitoring time profiling. Tests were conducted on a Lenovo ThinkStation P910 machine with Intel Xeon @ 2.3 GHz (28 cores) and 512 GB RAM.

### 6.2 Analysis of WGBS data

WGBS data of 27 different tissues was extracted from ENCODE (last download date: May of 2019, metadata on Table S2). Only tissue samples in biosample classification were selected. placenta and spinal cord tissues were discarded from the final dataset, and the remaining tissues were classified on endoderm, ectoderm or mesoderm germ layers. The resulting dataset consists of a matrix of 59 samples and 58 million CpG loci.

30 of the samples used QIAGEN DNeasy Blood & Tissue Kit as library extraction method, and nearly 3 million CpG loci were not included in the analysis. Those CpG loci, whose methylation level is 0 in those samples, and greater than zero in the rest, were removed. This filtering reduced the total amount of CpG loci to 55 million.

CpG loci with statistical significant differential methylation in the three germ layers were selected using our parallelized implementation of the KW test from scipy 1.3.1. CpG loci with a *p*-value smaller than 2· 10^−6^ were considered statistical significant and selected.

The chromosomal position of the statistical significant CpG loci, their nearest gene, their *p*-values and mean DNA methylation per layer are provided as a supplementary tabulated file.

In order to visualize the statistical significant CpG loci, clustermaps with cosine metric and average linkage methods were produced.

In order to assign the nearest gene to each CpG locus, gene boundaries were extracted as a bed file from UCSC table browser (hg38, Genes and Gene Predictions, knownCanonical options), and mappings were performed using bedtools 2.27.1 closest function.

Genes with more than 10 CpG loci in the matrix of CpG loci with statistical significant *p*-values were selected. Then, heatmaps of DNA methylation at the vicinity of each gene were used to represent methylation patterns upstream and downstream of the gene. CpG loci are represented upstream and downstream of the gene up to the farthest statistically significant CpG, plus CpGs within 5000 bases more. Then, the heatmap from each gene was manually inspected to find relevant methylation patterns upstream or downstream the gene, to do a subselection of the most representative cases.

### 6.3 Computation time analysis of WGBS data

To test the time efficiency of computation of KW test, we applied the test on the whole 55 million CpG loci dataset using 1, 2, 4, 8, 10, 15, 20 and 25 cores. Each measurement was repeated 5 times. For each case, two computation times were obtained: (1) the complete computation time, which accounts for the reading of the matrix, the *p*-value calculation and the writing of the results, and (2) reduced computation time, which accounts only for the *p*-value calculation, i.e. the parallelized part of the code.

In order to assess the efficiency of parallelization, we *log*_2_ transformed the number of cores and the reduced computation processing times, and adjusted the data to a linear regression. The slope represents the parallelization scaling which, in an ideal case, should be 1, that is, a linear scaling.

## Supporting information

Supplemental table, significant CpGs

## 7 Acknowledgments

This work have been supported by grants DFG113/18 from Diputación Foral de Gipuzkoa, Spain, Ministry of Economy and Competitiveness, Spain, MINECO grant BFU2016-77987-P, Basque Government Predoctoral Grant PRE_2018_1_0008, Spain, and Instituto de Salud Carlos III (AC17/00012) co-funded by the European Union (Era-cosysmed/H2020 Grant Agreement No. 643271).

The authors would like to thank Daniela Gerovska for fruitful discussion and comment during the preparation of this manuscript.

---

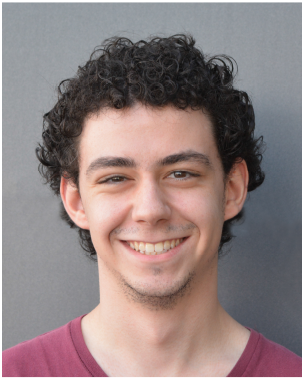

Alex M. Ascensión Graduated with honors of Biochemistry and Molecular Biology degree, ML gree, University of the Basque Country, Spain (2017). MSc in Bioinformatics, Autonomous University of Barcelona, Spain (2018). He is currently completing a Degree in Mathematics as a part-time student at the National University of Distance Education, Spain, to complement his biological and computational knowledge with a solid mathematics basis. Currently doing his PhD in Computational Biology with Basque Government grant. He is currently working at Biodonostia Health Research Institute, Spain, at the Computational Biology group (leadered by Marcos J. Araúzo-Bravo) alongside with the Tissue Engineering group (leadered by Ander Izeta). His research is focused on several topics, such as epigenomic signaling or singlecell studies. He is interested in the development and application of computational techniques and mathematical methods to his topics of research. He has more than 3 years of experience in the computational field, his main programming languages are Python and R, as well as Matlab, Perl or C++ to a lesser extent. He is a member of the scientific committee of the European Project at the Eracosysmed JTC-2 (H2020) call 4D-Healing: Data-Driven Drug Discovery for Wound Healing. His main functions within the committee are the development of the web page and the computational analysis of the generated single-cell RNA-seq data.

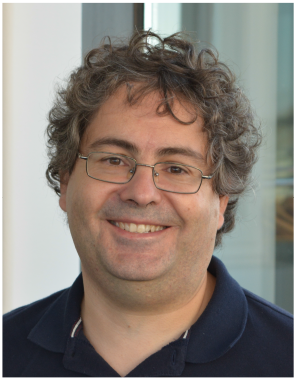

Marcos J. Araúzo-Bravo Graduated as an Electronic and Control Engineer, University of Valladolid, Spain (1996). In 2001 he earned a Ph.D. in industrial technologies from the University of Cartagena, developing neuro-fuzzy algorithms for monitoring penicillin production. From 1998 to 2004 he was an Associate Professor in electrical engineering at Burgos University. In 2000 he received a scholarship from the Japanese Ministry of Education to work in the field of Metabolic Engineering at Kyushu Institute of Technology, Japan. In 2002 he earned a Ph.D. in Information Technology and Biotechnology from the Kyushu Institute of Technology, Iizuka, Japan. From 2004 to 2006 he was a Japan-Society-for-the-Promotion-of-Science postdoctoral research fellow at the Kyushu Institute of Technology, where he worked on the synergetic control of genetic networks through transcriptional regulators. From 2006 to 2014 he led the laboratory of Computational Biology and Bioinformatics at the Max Planck Institute for Molecular Biomedicine in Münster, Germany, developing tools for deciphering cellular reprogramming, methods to study transcription regulation, and algorithms for high-throughput data analysis. Since 2014 he is an Ikerbasque Research Professor, head of the group of Computational Biology and Systems Biomedicine and head of the Computational Biomedicine Data Analysis Platform at the Biodonostia Health Research Institute, San Sebastián, Spain. He develops Big Data approaches for integrating omics, image and clinical history data to study the interaction of biological networks in terms of their topology, dynamics, and perturbations to interpret complex biological systems associated with neurodegenerative diseases, cancer, aging, stem cells and regenerative medicine. His vast experience analyzing and deriving models from omics data produced more than 130 publications, some of them very high impact such as Science, Nature, Nature, Cell. He is the coordinator of the European Project at the Eracosysmed JTC-2 (H2020) call 4D-Healing: Data-Driven Drug Discovery for Wound Healing.

