## Supplemental table, significant CpGs for "BigMPI4py: Python module for parallelization of Big Data objects"

March 10, 2020

#### Supplementary tables and figures

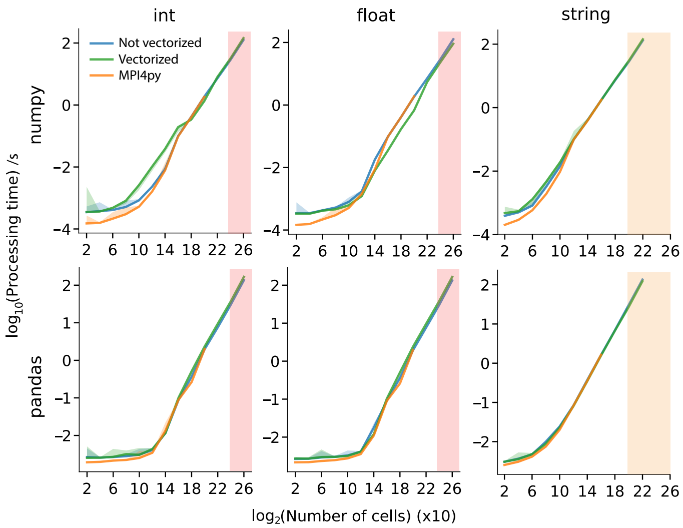


Figure S1: **Gathering times for numpy arrays and pandas dataframes in three data types.** The number of cells that MPI4py is not able to process due to *OverflowError* is shadowed in pink for int and float data types, and in orange for string data type. Color filling between lines represents the $10^{th}$ and $90^{th}$ percentiles of the data.

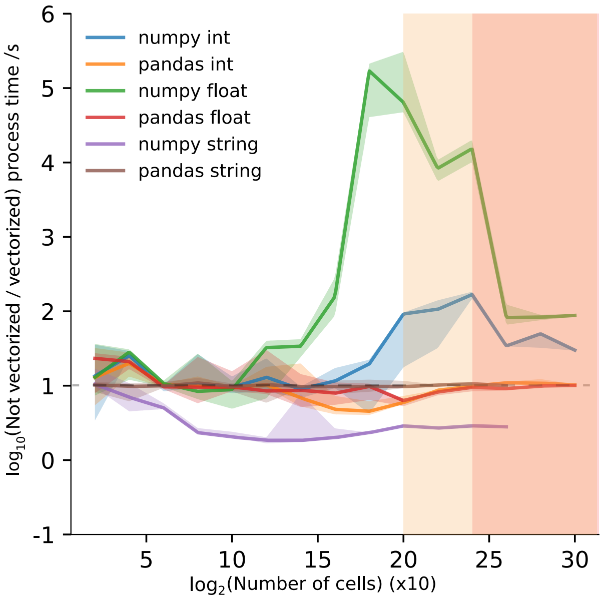


Figure S2: **Scattering time ratio between non-vectorized and vectorized approaches.** The number of cells that MPI4py is not able to process due to *OverflowError* is shadowed in pink for int and float data types, and in orange for string data type. Color filling between lines represents the $10^{th}$ and $90^{th}$ percentiles of the data.

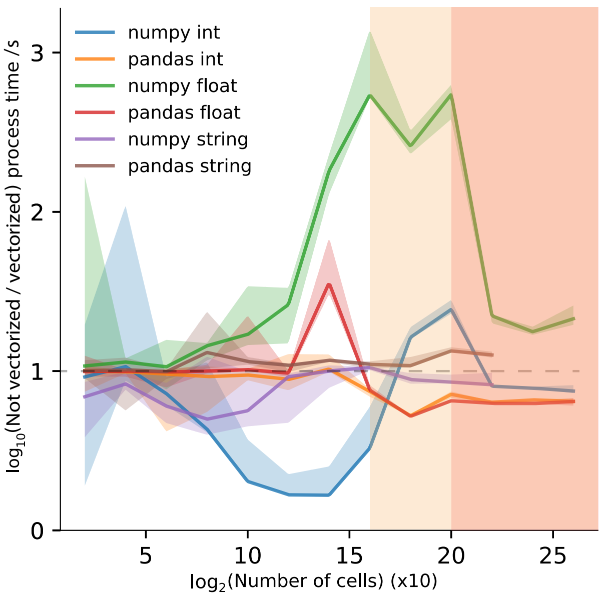


Figure S3: **Gathering time ratio between non-vectorized and vectorized approaches.** The number of cells that MPI4py is not able to process due to *OverflowError* is shadowed in pink for int and float data types, and in orange for string data type. Color filling between lines represents to the $10^{th}$ and $90^{th}$ percentiles of the data.

Table S1: Selection of genes from the list of statistically significant different methylated *loci* with ion patterns associated to germ layers.

| Ectoderm + mesoderm | Endoderm |
|---|---|
| AL121722.1 | AC018521.1 |
| COX7A1 | AC092153.1 |
| DDR2 | AC243965.1 |
| FAM198B | BCL2 |
| IFFO1 | BHLHE41 |
| LINC00261 | C1S |
| LINC01571 | CDH1 |
| LINC02325 | DNMBP-AS1 |
| MNX1 | DNMBP |
| PRKACA | ELF3 |
| PRRT1B | ERVE-1 |
| SSH1 | KLF5 |
| ST3GAL4 | LPCAT3 |
| TCF4 | MIR200B |
| TTC6 | MIR200CHG |
| | PBX1 |
| | PLLP |
| | PRR15L |
| | RBM47 |
| | TMEM51 |

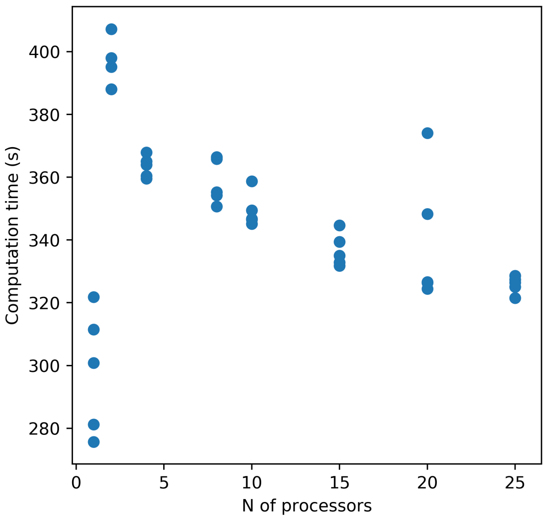


Figure S4: **Base computation times for KW test on WGBS data.** Base computation time is defined as the time associated to non-parallelizable actions like reading or writing of data. Computation times have been replicated 5 times for each processor number option. Base computation times have a relatively narrow time window, despite de deviation associated to number of processors, and compared to times associated to parallel processing

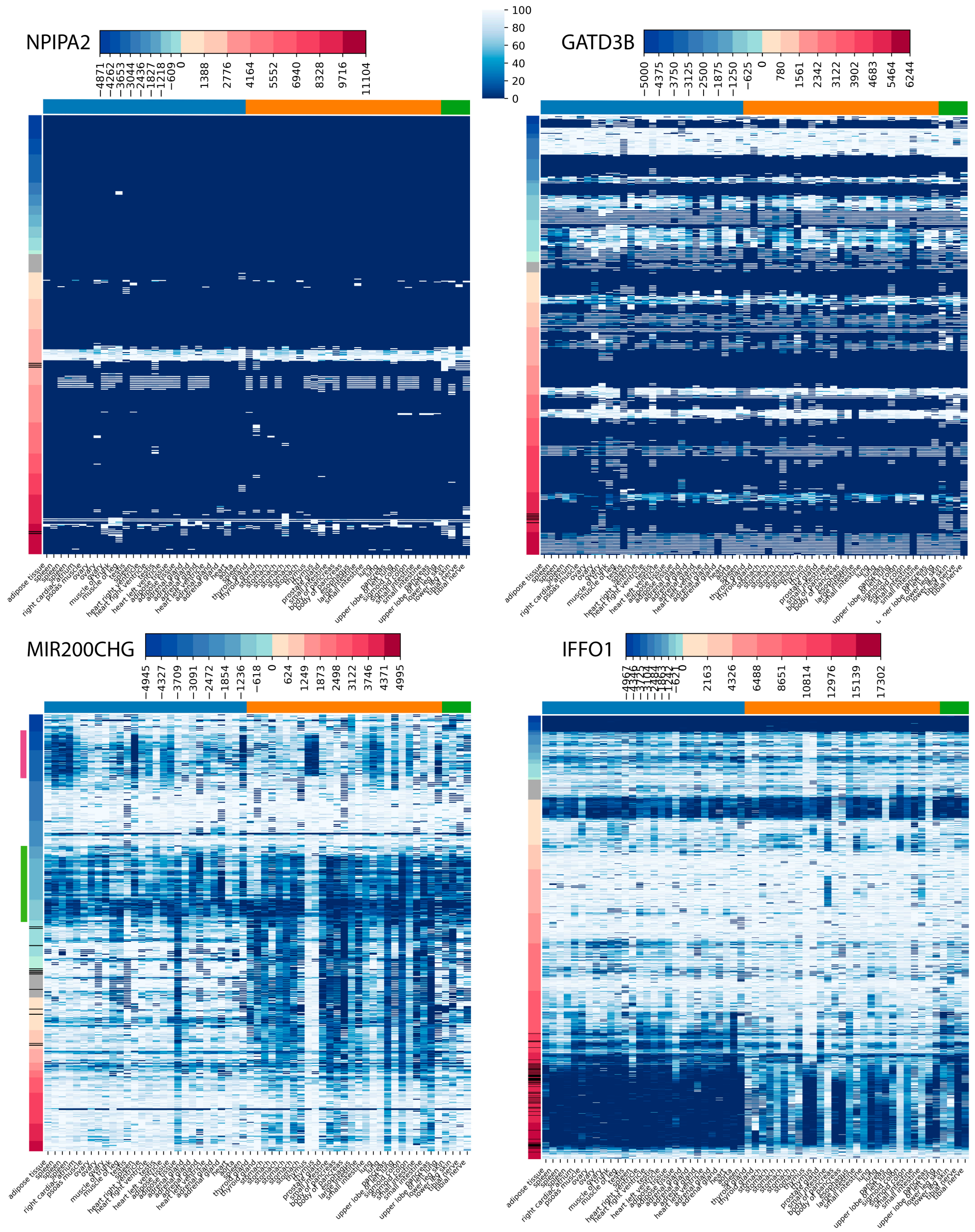


Figure S5: **Heatmaps of methylation around *NPIPA2*, *GATD3B*, *MIR200CHG* and *IFFO1* genes.** Heatmap columns are different manually-labelled tissue types (orange: endoderm, blue: mesoderm, green: ectoderm), and rows are CpG *loci*. Coloured labels in rows represent the distance from the gene, in bases. Gray label represents CpG *loci* within the gene, blue labels represent positions upstream the gene, and red labels represent positions downstream the gene. Black marks in row labels represent positions with $p$-value $< 2 \cdot 10^{-6}$. For *MIR200CHG* gene the region with high DNA methylation in thymus, and slight methylation level in spleen is marked in green on the left; and the upstream region showing a strong demethylation of both tissues is shown in pink. The color bar in the top right codifies the CpG methylation in percentage. Higher methylation corresponds to whiter color.

Table S2: Description of ENCODE sample metadata.

| File accession | Biosample term name | Germ layer | Library extraction method | Size |
|---|---|---|---|---|
| ENCFF923CZC | large intestine | endoderm | | 639828675 |

| | | | | |
|---|---|---|---|---|
| ENCFF913ZNZ | adrenal gland | mesoderm | | 666396871 |
| ENCFF844EFX | stomach | endoderm | | 686198683 |
| ENCFF843SYR | tibial nerve | ectoderm | | 691993425 |
| ENCFF842MHJ | upper lobe of left lung | endoderm | | 671508093 |
| ENCFF831OYO | heart right ventricle | mesoderm | QIAGEN DNeasy Blood & Tissue Kit | 658134692 |
| ENCFF826PSA | adrenal gland | mesoderm | | 644150581 |
| ENCFF811QOG | stomach | endoderm | | 685147881 |
| ENCFF763RUE | pancreas | endoderm | QIAGEN DNeasy Blood & Tissue Kit | 734876240 |
| ENCFF748MTS | body of pancreas | endoderm | | 666588523 |
| ENCFF733EFJ | upper lobe of left lung | endoderm | | 675759526 |
| ENCFF730NQT | spleen | mesoderm | QIAGEN DNeasy Blood & Tissue Kit | 663517254 |
| ENCFF715DMX | testis | mesoderm | | 639650311 |
| ENCFF699RBP | body of pancreas | endoderm | | 631023722 |
| ENCFF699KTW | tibial nerve | ectoderm | | 686090388 |
| ENCFF684JHX | heart left ventricle | mesoderm | QIAGEN DNeasy Blood & Tissue Kit | 712653475 |
| ENCFF645AZF | muscle of trunk | mesoderm | | 688503164 |
| ENCFF638QVP | testis | mesoderm | | 692606437 |
| ENCFF625GVK | esophagus | endoderm | QIAGEN DNeasy Blood & Tissue Kit | 732064586 |
| ENCFF618WAT | adrenal gland | mesoderm | | 672354523 |
| ENCFF588ETU | muscle of leg | mesoderm | | 663027933 |
| ENCFF560SMW | heart | mesoderm | | 696737981 |
| ENCFF553HJV | aorta | mesoderm | QIAGEN DNeasy Blood & Tissue Kit | 650626532 |
| ENCFF550FZT | spleen | mesoderm | | 688273627 |
| ENCFF536RSX | heart left ventricle | mesoderm | QIAGEN DNeasy Blood & Tissue Kit | 732733386 |
| ENCFF534RNT | stomach | endoderm | QIAGEN DNeasy Blood & Tissue Kit | 659322648 |
| ENCFF526PFA | spleen | mesoderm | QIAGEN DNeasy Blood & Tissue Kit | 731972903 |
| ENCFF521DHD | small intestine | endoderm | | 616061802 |
| ENCFF513ITC | heart right ventricle | mesoderm | QIAGEN DNeasy Blood & Tissue Kit | 708249770 |
| ENCFF500DKA | pancreas | endoderm | QIAGEN DNeasy Blood & Tissue Kit | 653548206 |
| ENCFF497YOO | stomach | endoderm | QIAGEN DNeasy Blood & Tissue Kit | 709076242 |
| ENCFF497IYX | thyroid gland | endoderm | | 674810141 |
| ENCFF489CEV | stomach | endoderm | | 685734577 |
| ENCFF477GKI | adipose tissue | mesoderm | QIAGEN DNeasy Blood & Tissue Kit | 656844077 |
| ENCFF477AUC | lung | endoderm | QIAGEN DNeasy Blood & Tissue Kit | 645181452 |
| ENCFF455TQO | sigmoid colon | endoderm | QIAGEN DNeasy Blood & Tissue Kit | 727685810 |
| ENCFF435SPL | stomach | endoderm | QIAGEN DNeasy Blood & Tissue Kit | 663904217 |
| ENCFF392XPZ | thymus | endoderm | | 639152431 |
| ENCFF333OHK | spleen | mesoderm | | 678390529 |
| ENCFF318AMC | adipose tissue | mesoderm | QIAGEN DNeasy Blood & Tissue Kit | 714923500 |
| ENCFF303ZGP | ovary | mesoderm | | 687213122 |
| ENCFF266NGW | small intestine | endoderm | QIAGEN DNeasy Blood & Tissue Kit | 646656686 |
| ENCFF247ILV | ovary | mesoderm | QIAGEN DNeasy Blood & Tissue Kit | 714367475 |
| ENCFF241AQC | small intestine | endoderm | QIAGEN DNeasy Blood & Tissue Kit | 733852691 |
| ENCFF223LJW | thyroid gland | endoderm | | 672601274 |
| ENCFF219GCQ | lower leg skin | ectoderm | | 689355695 |
| ENCFF216DJL | adrenal gland | mesoderm | QIAGEN DNeasy Blood & Tissue Kit | 708311426 |
| ENCFF210XTE | adrenal gland | mesoderm | QIAGEN DNeasy Blood & Tissue Kit | 662267042 |
| ENCFF200MJQ | spleen | mesoderm | QIAGEN DNeasy Blood & Tissue Kit | 652870215 |
| ENCFF189WPY | ovary | mesoderm | | 686846640 |
| ENCFF168HTX | thymus | endoderm | QIAGEN DNeasy Blood & Tissue Kit | 693624498 |
| ENCFF157POM | sigmoid colon | endoderm | QIAGEN DNeasy Blood & Tissue Kit | 735717264 |
| ENCFF122LEF | small intestine | endoderm | QIAGEN DNeasy Blood & Tissue Kit | 649925601 |
| ENCFF121ZES | psoas muscle | mesoderm | QIAGEN DNeasy Blood & Tissue Kit | 649650845 |
| ENCFF121VIX | lower leg skin | ectoderm | | 692313819 |

| | | | | |
|---|---|---|---|---|
| ENCFF110AZO | right cardiac atrium | mesoderm | QIAGEN DNeasy Blood & Tissue Kit | 713431488 |
| ENCFF103DNU | adipose tissue | mesoderm | QIAGEN DNeasy Blood & Tissue Kit | 652018078 |
| ENCFF039JFT | lung | endoderm | QIAGEN DNeasy Blood & Tissue Kit | 725135542 |
| ENCFF027KTR | prostate gland | endoderm | | 674251116 |

### Supplementary algorithms

**Algorithm S1** Strategy I for scattering

```
procedure STRATEGY_I_SCATTER(A, idx_A, L, n)
    if rank == root then
        k ← 0
        idx_k ← []
        scatter_A ← []
        for i in 0:len(idx_A) do
            k ← max(k, int(size(A[idx_A[i:i+1]]) / L) + 1)
        end for
        for i in 0:len(idx_A) - 1 do
            APPEND(idx_k, int(z) for z in np.linspace(idx_A[i], idx_A[i + 1], k + 1)][:-1])
        end for
        APPEND(idx_k, len(A))
        for i in 0:len(idx_k) - 1 do
            APPEND(scatter_A, A[idx_k[i]:idx_k[i + 1]])
        end for
    end if
    k ← SCATTER(k)
    scatter_object ← [None] * k
    for ki in 0:k do
        scatter_object[ki] ← SCATTER([scatter_A[i] for i in range(0, k * n, k)])
        for i in 0:k * n:k do                                  ▷ Deleting to free memory
            del scatter_A[i]
        end for
    end for
    return MERGE(scatter_object)
end procedure
```

**Algorithm S2** Strategy I for gathering

```
procedure STRATEGY_I_GATHER(A, L, n)
    k ← 0
    divide_A ← []
    for i in 0:len(idx_A) do
        k ← int(size(A) / L) + 1
    end for
    k ← max(ALLGATHER(k))
    idx_k ← [int(z) for z in np.linspace(0, len(A), k + 1)]
    for i in 0:len(idx_k) - 1 do
        APPEND(divide_A, A[idx_k[i]:idx_k[i + 1]])
    end for
    if rank == root then
        gather_list ← [None] * (n * k)
    end if
    for ki in 0:k do
        gather_ki ← GATHER(divide_A[ki])
        for ni in 0:n do
            gather_list[ki + ni * k] = gather_ki[i]
        end for
    end for
    return MERGE(gather_list)
end procedure
```

**Algorithm S3** Strategy II for scattering

```
procedure STRATEGY_II_SCATTER(A, idx_A, L, n)
    k ← 0
    for i in A do
        k ← max(k, int(size(i) / L) + 1)
    end for
     scatter_list ← [[[] for ki in k] for ni in n]
    for ni in n do
        A_ni ← A[idx_A[ni]:idx_A[ni+1]
        A_ni_k ← []
        for obj in A_ni do
            idx_k ← [int(i) for i in LINSPACE(0, len(obj), k+1)]
            obj_k ← [obj[idx_k[i]:idx_k[i+1] for i in 0:k]
            APPEND(A_ni_k, obj_k)
        end for
        scatter_list[ni] ← A_ni_k
    end for
     merge ← []
    for ki in 0:k do
        if rank == root then
            merge_ki ← []
            for sc_list_ni in sc_list do
                x ← []
                for l in 0:len(sc_list_ni) do
                    APPEND(x, sc_list_ni[l][ki])
                end for
                APPEND(merge_ki, x)
            end for
        end if
        merge_ki ← SCATTER(merge_ki)
        APPEND(merge, merge_ki)
    end for
    return_list ← []
    for l in 0:len(merge[0]) do
        return_ki ← [merge[ki][l] for ki in 0:k]
        APPEND(return_list, MERGE(return_ki))
    end for
    return return_list
end procedure
```

**Algorithm S4** Strategy II for gathering

```
procedure STRATEGY_II_GATHER(A, L, n)
    k ← 0
    for i in A do
        k ← max(k, int(size(i) / L) + 1)
    end for
    k ← max(ALLGATHER(k))
    max_len_A ← len(A)
    max_len_A ← max(ALLGATHER(max_len_A))
    A ← A + [[] * (max_len_A - len(A))]
    gather_list ← [None] * (n * max_len_A)
    for i in 0:len(max_size_list) do
        cut_idx ← [int(x) for x in LINSPACE(0, len(i), k+1)]
        return_i ← []
        for ki in range(k) do
            x ← A[i][cut_idx[ki]: cut_idx[ki+1]
            gather_obj ← GATHER(x)
            APPEND(return_i, gather_obj)
        end for
        if return_i[0] then
            for ni in 0:n do
                list_return_i ← [return_i[ki][ni] for ki in range(k)]
                merged_i ← MERGE(list_return_i)
                if len(merged_i) > 0 then
                    gather_list[i + ni*max_len_A] ← merged_i
                end if
            end for
        end if
    end for
    return gather_list
end procedure
```

**Algorithm S5** bcast() algorithm

```
procedure BCAST(A, L, n)
    mem ← GET_MEMORY()
    size_A ← GET_SIZE(A)
    for n_i in 0:n do
        bcast_list ← [None] * n
        bcast_list[n_i] ← A
        bcast_obj ← SCATTER(bcast_list, L)
        if rank == n_i then
            return_obj ← bcast_obj
        end if
    end for
    return return_obj
end procedure
```

**Algorithm S6** Object gathering and allgathering

```
arr = np.random.rand(2 ** 26, 10)
gather_arr = BM.gather(arr, comm)
allgather_arr = BM.allgather(arr, comm)
```

**Algorithm S7** Object point-to-point communication

```
arr = np.random.rand(2 ** 26, 10)
sendrecv_arr = BM.sendrecv(arr, comm,
dest = 2)
```

**Algorithm S8** Example of MPI launch

```
mpirun -np 4 python ~/mycode.py
```

### Supporting information

**Example of list scattering with Strategy I**

$A = [a_1, a_2, a_3, \cdots, a_9]$ is distributed into $n = 5$ cores, thus it is divided into 5 *chunks*: $A_1 = [a_1, a_2]$, $A_2 = [a_3, a_4]$, $\cdots$, $A_5 = [a_9]$. If $k_I = 2$, then the distribution of *subchunks* is $[[a_1], [a_2], [a_3], [a_4], \cdots, [a_9], []]$, i.e., $A_{1,1} = a_1, A_{1,2} = a_2, A_{1,3} = a_3, \cdots, A_{5,1} = a_9, A_{5,2} = []$. After scattering, each core receives the list $[A_{i,1}, A_{i,2}]$; for core 3 this list is $[[a_5], [a_6]]$, and for core 5 it is $[[a_9], []]$. After merging, core 3 has $[a_5, a_6]$ and core 5 has $[a_9]$.

**Example of gathering with Strategy II**

The first core has the list $A_1 = [a_1, a_2, a_3]$, and the second core has the list $A_2 = [a_4, a_5]$. We obtain a $k_{II}$ value of 2 for the second core, and of 4 for the first core, thus $k_{II} = 4$. $A_2$ is filled to $[a_4, a_5, []]$, and `gather_list` is [`None`, `None`, `None`, `None`, `None`, `None`]. BigMPI4py iterates for `obj` in `0:3` and, for each iteration, gathers the $i$-th value of $A_1$ and $A_2$. The first iteration takes $a_1$ and $a_4$ respectively. Next, iterating for $k_i$, `x` $= a_{1,1}$ and $a_{4,1}$ for each core, and `gather_obj` $= [a_{1,1}, a_{4,1}]$. After all $k_i$ iterations, the list of objects $[[a_{1,1}, a_{4,1}], [a_{1,2}, a_{4,2}], \cdots, [a_{1,4}, a_{4,4}]]$ is created. Then, iterating through $n$, $[a_{1,1}, \cdots, a_{1,4}]$ and $[a_{4,1}, \cdots, a_{4,4}]$ are merged to $a_1$ and $a_4$, and `gather_list` is filled to be [$a_1$, `None`, `None`, $a_4$, `None`, `None`]. After all iterations, `gather_list` is [$a_1$, $a_2$, $a_3$, $a_4$, $a_5$, `None`], the `None` element is removed, and the list is returned.
